# Alka(e)nes contribute to membrane lipid homeostasis and resilience of photosynthesis to high light in cyanobacteria

**DOI:** 10.1101/2023.12.11.571064

**Authors:** Rui Miao, Bertrand Légeret, Stéphan Cuine, Adrien Burlacot, Peter Lindblad, Yonghua Li-Beisson, Frédéric Beisson, Gilles Peltier

## Abstract

Alka(e)nes are produced by many living organisms and exhibit diverse physiological roles, reflecting a high functional versatility. Alka(e)nes serve as water proof wax in plants, communicating pheromones for insects, and microbial signaling molecules in some bacteria. Although alka(e)nes have been found in cyanobacteria and algal chloroplasts, a possible role in photosynthesis and chloroplast function remains elusive. In this study, we investigated the consequences of the absence of alka(e)nes on membrane lipid remodeling and photosynthesis using the cyanobacteria *Synechocystis* PCC6803 as a model organism. By following the dynamics of membrane lipids and the photosynthetic performance in strains defected and altered in alka(e)ne biosynthesis, we show that a profound remodeling of the membrane lipidome and carotenoid content occur in the absence of alka(e)nes, including a decrease in the membrane carotenoid content, a decrease in some digalactosyldiacylglycerol (DGDG) species and a parallel increase in monogalactosyldiacylglycerol (MGDG) species. Under high light, this effect is accompanied in alka(e)ne deficient strains by a higher susceptibility of photosynthesis and growth, the effect being reversed by expressing an algal photoenzyme producing alka(e)nes from fatty acids. We conclude that alka(e)nes play a crucial role in maintaining lipid homeostasis of photosynthetic membranes, thereby contributing to the proper functioning of photosynthesis, particularly under elevated light intensities.

**Significance statement:** We used cyanobacteria as a model organism to explore the role of alka(e)nes related to photosynthesis. Our findings reveal that the absence of alka(e)nes induces alterations in the composition of membrane lipids and carotenoid content, resulting in an increased susceptibility of photosynthesis. By introducing a fatty acid photodecarboxylase to produce alkanes, we could reverse these effects, highlighting the critical role of alka(e)nes in maintaining lipid balance in photosynthetic membranes and ensuring efficient photosynthesis. Uncovering the physiological role of alka(e)nes provides insights to a better understanding of the widespread presence of genes encoding alka(e)nes-synthesizing enzymes in cyanobacteria and microalgae, organisms of major ecological and evolutionary importance in the global CO_2_ assimilation.

## Introduction

Alkanes (saturated) and alkenes (unsaturated), both classes of acyclic hydrocarbons and the principal components of fossil fuels, are ubiquitous in our daily lives, serving diverse functions and applications. Remarkably, alka(e)nes are not confined to the realm of fossil fuels; numerous organisms across the biological kingdom also synthesize them, each with distinct carbon chain lengths, cellular locations and physiological functions. For instance, insects primarily synthesize very-long-chain linear alka(e)nes (*n*-alka(e)nes) through a process that involves fatty acid elongation, reduction to aldehyde and subsequent oxidative decarbonylation (1–3). These *n*-alka(e)nes serve essential roles in preventing desiccation and act as potent sensory neuron activators during defensive and reproductive behaviors (3, 4). Plants also synthesize *n*-alka(e)nes from fatty acids, with cuticular waxes enriched in very-long-chain alkanes providing protection against desiccation and environmental stressors like UV radiation and pathogens (5, 6). Among unicellular organisms, some non-photosynthetic bacteria have the ability to naturally produce alka(e)nes, primarily via the decarbonylation of fatty aldehydes or decarboxylation of fatty acids (7, 8). As photosynthetic microorganisms, microalgae also harbor a distinctive pathway for alka(e)ne synthesis (9). The discovery of fatty acid photodecarboxylase (FAP) in green microalgae (10) has enriched our repertoire of alka(e)ne-forming enzymes because of the unique photocatalytic process of FAP, which converts free fatty acids (FFA) to alka(e)nes using light in the range 350-530 nm (11, 12). FAP belongs to an algal-specific clade of glucose methanol choline (GMC) oxidoreductase family and phylogenetic analysis showed that this photoenzyme is conserved in photosynthesis-retained algal lineages (13). In cyanobacteria (oxygenic photosynthetic bacteria), alka(e)ne biosynthesis is also a well-established phenomenon and has been subsequently optimized by metabolic engineering (14–17). Cyanobacteria, are known to harbor two mutually exclusive alka(e)ne synthesis pathways: the acyl-ACP reductase (Aar) and aldehyde deformylating oxygenase (Ado) pathway (18) and the olefin synthase (Ols) pathway (19). In a survey of 73 cyanobacterial strains spanning a broad phylogenetic range, it was found that the vast majority possessed the Aar-Ado pathway and only 12 strains, two-third of which were from a single clade, possessed the Ols pathway (20). It is interesting to note that while the alkanes and alkenes synthesized by cyanobacteria and microalgae represent only a few percent at most of their total fatty acids, on a global scale this synthesis is not anecdotal, as an oceanic cycle of long-chain alkanes produced by marine cyanobacteria (and also probably microalgae) has been identified and characterized (21, 22).

The diverse alka(e)ne biosynthesis pathways in nature highlight the intricate evolutionary adaptations of organisms to their ecological niches and the multifaceted roles of alka(e)nes in various biological contexts. Notably, orthologous genes to *aar* and *ado* have been identified exclusively in cyanobacteria so far, implying a potential connection to a photoautotrophic lifestyle (23). Photosynthesis, a delicate and complex process, relies on the complex structure and fluidity of cellular membranes, particularly the thylakoid membrane where light-dependent reactions occur. In the model microalga *Chlamydomonas reinhardtii*, FAP and its major product 7-heptadecene were found associated to the thylakoid membranes in chloroplast, and the study of a FAP knock-out mutant suggested a role for this alkene in photosynthesis under cold and varying light conditions but the exact molecular mechanism remains unknown (13). The physiological function of the native intracellular alka(e)nes has only recently started to be investigated in cyanobacteria, and studies on strains with the *aar/ado* pathway knocked out revealed various deficiencies. In *Synechococcus* PCC 7002 and *Synechocystis* PCC 6803 alka(e)nes-deficient strains, reduced growth and impaired cell division were observed (24). Without alka(e)nes, *Synechocystis* also showed poor growth and diminished photosynthetic cyclic electron flow at low temperatures (25), and *Synechococcus elongatus* PCC 7942 exhibited decreased salt tolerance (26). In addition, molecular dynamic simulation was employed to investigate alka(e)nes organization in lipid membranes, suggesting that alka(e)nes localize in the middle part of the lipid bilayer, and may play a role in membranes by promoting flexibility and facilitating curvature (24).

In this study, we use *Synechocystis* PCC6803 as a model organism to bridge the significant knowledge gap concerning the relationship between alka(e)nes and photosynthesis, as well as to explore the potential underlying molecular mechanisms. By analyzing the lipidome dynamics and the activity of photosynthesis in response to changing light conditions, we conclude that alka(e)nes contribute to membrane lipid homeostasis and participate in the resilience of photosynthesis to high light.

## Results

### Absence of alka(e)nes affect cell ultrastructure

In *Synechocystis*, as in many other cyanobacterial strains, the acyl-ACP reductase encoding gene *aar* and the deformylating oxygenase encoding gene *ado* are transcribed in two neighboring operons (20, 23). Two transcription start sites (TSS) have been identified at the 5’-end of the *ado* gene, which may imply the potential transcription of multiple isoforms under specific conditions (23). Considering the regulatory implications of such complex transcriptional organization, we generated three alka(e)ne-deficient *Synechocystis* strains, ΔAdo, ΔAar, and ΔAdoAar. The deformylating oxygenase Ado uses an external reducing system capable of supplying four electrons for each catalytic cycle, so alka(e)ne synthesis has been considered as an electron sink to dissipate exceed energy in specific conditions (23, 27). Therefore, to attribute observed phenotypes to alkanes by avoiding potential effects of electron consumption and metabolic intermediates, we used the photoenzyme fatty acid photodecarboxylase (FAP) from *Chlorella variabilis* (10) to functionally complement strains deficient in alka(e)nes. Unlike the Aar-Ado pathway, which utilizes acyl-ACPs as substrates and produces fatty aldehydes as intermediates, the FAP enzyme catalyzes the decarboxylation of a free fatty acid in a single step, requiring light in the range of 350-530 nm.

FAP and a chloramphenicol resistance gene weres integrated into the genome of each alka(e)ne-deficient strain at the *slr1993* locus, resulting in strains ΔAdo-FAP, ΔAar-FAP, and ΔAdoAar-FAP. *Slr1993* encodes the acetyl-CoA acetyltransferase PhaA, and deleting PhaA may eliminate resource competition between PHB synthesis and alka(e)nes synthesis, which may ensure a decent level of alkane production from FAP. Concurrently, we generated corresponding control strains denoted as ΔAdo-FAP control, ΔAar-FAP control, and ΔAdoAar-FAP control by integrating a chloramphenicol resistance gene at *slr1993* locus. As previously reported (18, 20), the major alka(e)ne produced by *Synechocystis* was found to be *n*-heptadecane (C17 *n*-alkane), but minor amounts of *n*-pentadecane (C15 *n*-alkane) and two heptadecene isomers (C17 mono-alkene) were also detected (Fig. 1A). As anticipated, no alka(e)nes were detectable in strains ΔAdo, ΔAar, and ΔAdoAar. Complemented strains expressing FAP produced the same type of alka(e)nes as WT strain, while at lower amounts particularly for the more abundant species heptadecane (Fig. 1A).

**Fig. 1.**
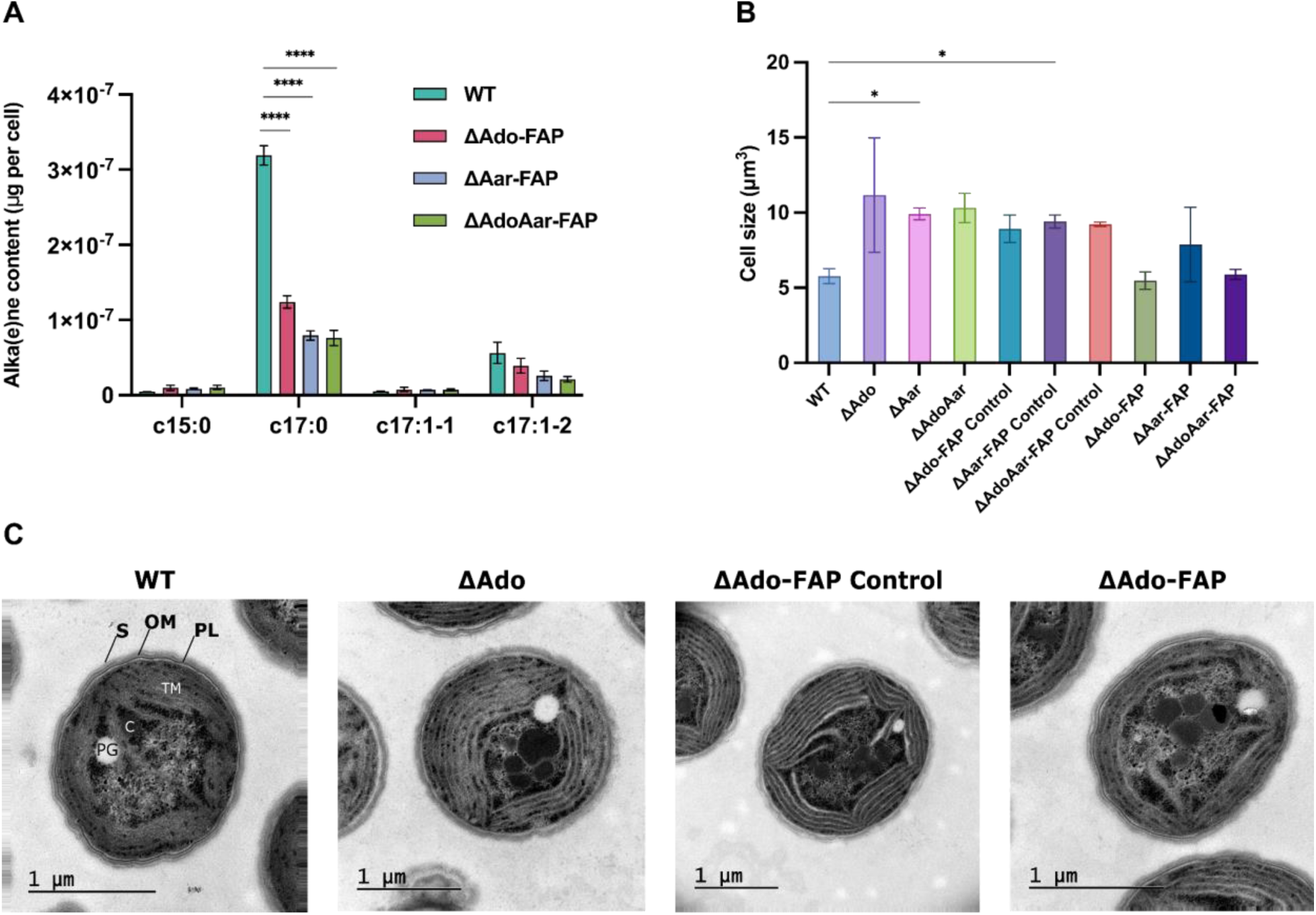
Alka(e)nes production, cell size, and cell morphology in *Synechocystis* PCC6803 strains cultivated under 30 μmol photon m^−2^ s^−1^. **A.** Alka(e)nes production in wild-type and all complemented strains. Alka(e)nes were quantified from mid-log phase (OD_730_=0.6) cultures using transmethylation of cells and GC-MS analysis. Alka(e)ne production was normalized to cell number. Results represent the mean of 3 biological replicates; error bars represent standard deviation. Two-way ANOVA was used to determine statistical significance. **B.** Cell size comparison among all strains. All cells were grown to mid-log phase (OD_730_=0.6), and cell volume was measured using a Coulter counter. Results represent the mean of 3 biological replicates; error bars represent standard deviation. One-way ANOVA was used to determine significance. **C.** Transmission electron microscopy images. All cells were grown to OD_730_=0.3 then for room temperature fixation. TM: thylakoid membrane; **C:** carboxysome; PG: polyphosphate granule; PL: peptidoglycan layer; OM: outer membrane; S: S-layer.

In line with a previous study (24), we observed that the cell size of the alka(e)ne-deficient strains exhibited a significant increase, with dimensions nearly doubling that of both the WT strain and the complemented strains (Fig. 1B). To further gain information on subcellular changes, we examined the cells under transmission electron microscopy (TEM) which revealed a profound impact on the organization of thylakoid membranes in the absence of alka(e)nes (Fig. 1C). While the WT and complemented strains consistently exhibited between 4 and 5 thylakoid layers per cell, the alka(e)ne-deficient cells seemed to have 7 apparent layers of thylakoid membranes per cell (Fig. 1C). This transformation in thylakoid structure suggests a key role for alka(e)nes in maintaining membrane architecture, with potential repercussions on photosynthetic capacity.

Moreover, a closer inspection of the cell surface revealed pronounced differences. In alka(e)ne-deficient strains, the S-layer and peptidoglycan layer surrounding the cells appeared unusually blurred and disrupted, in stark contrast to the well-defined appearance observed in the WT and complemented strains. These results suggest that part of the hydrocarbons formed interact with cell surface components or influence cell wall formation.

### Alka(e)nes are critical for cell growth and photosynthesis under high light

We then aimed at determining whether alka(e)ne-deficiency affects photoautotrophic growth by performing growth tests on solid media under varying light intensities. For each strain, experiments were initiated with three different initial cell concentrations. While no growth difference was observed under low light (20 μmol photons m^-2^ s^-1^) between the different strains, a significant growth defect was observed in all alka(e)ne-deficient strains under medium light (50 μmol photons m^-2^ s^-1^), and a complete growth arrest under high light (200 μmol photons m^-2^ s^-1^) (Fig. 2A). Remarkably, this growth phenotype was fully restored in the complemented line despite a lower accumulation of alkanes as compared to WT strain (Fig. 1A). Because strains ΔAdo, ΔAar, and ΔAdoAar always showed similar phenotypes (Supplementary figure 1), further experiments were performed on one of them, ΔAdo, and its corresponding controls.

**Fig. 2.**
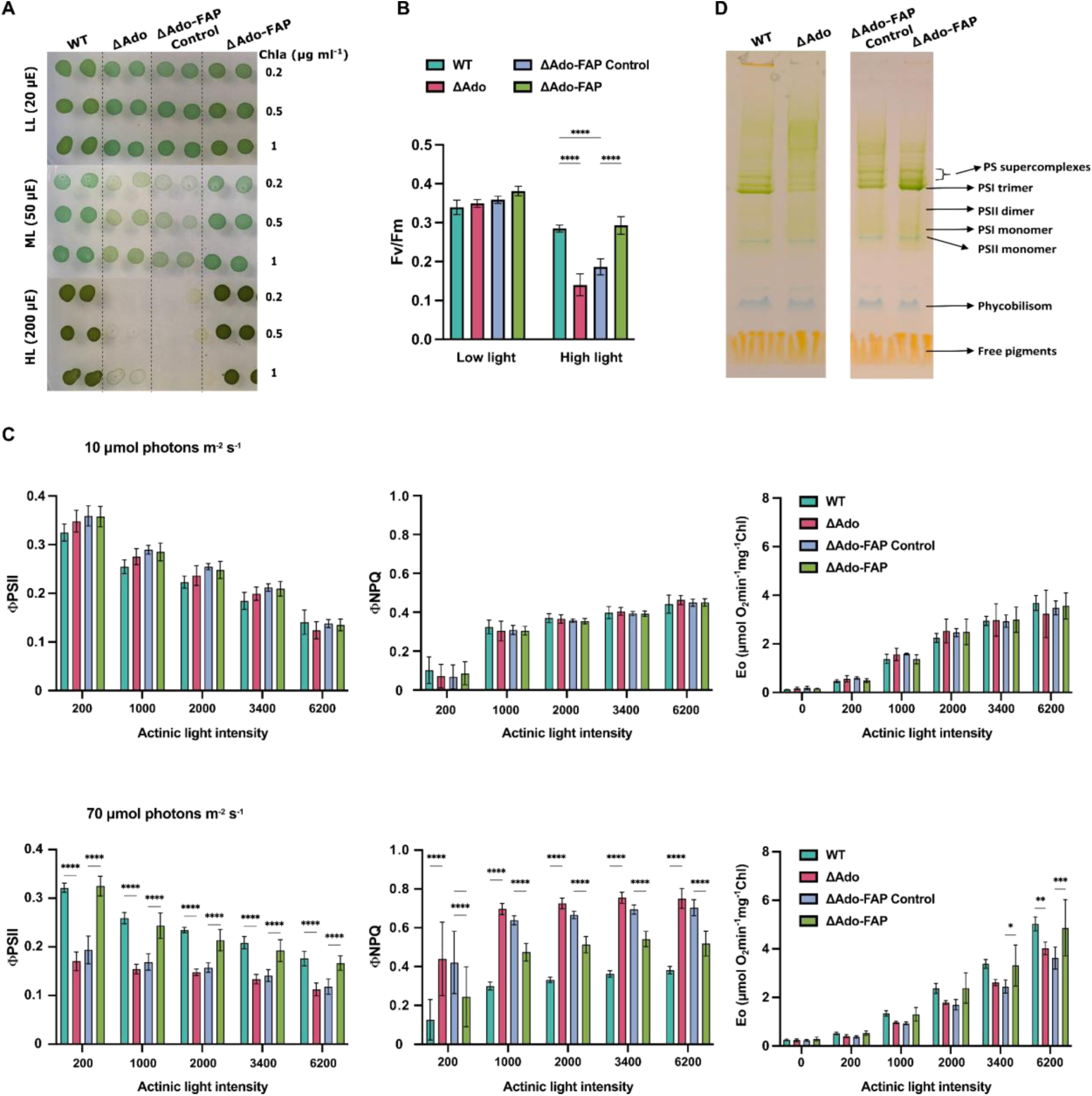
Alka(e)nes effects on cell growth test and photosynthetic capacity. **A.** Growth test on BG-11 agar plate under different light intensities. Biological duplicate and three different chlorophyll concentrations were used for each strain. (1) Wild-type (2) ΔAdo (3) ΔAdo-FAP Control (4) ΔAdo-FAP. **B.** The quantum yield (Fv/Fm) for dark-adapted cells was measured after 10 minutes of darkness adaptation. These cells had previously been cultivated under either low light or high light conditions. **C.** Simultaneous measurements of operating PSII quantum yield (ΦPSII), non-photochemical quenching yield (ΦNPQ), and Membrane-inlet-mass-spectrometry (MIMS) measured gross O_2_ production rates (Eo) on cultures grown under both low and high light conditions. All the measurements were done under increased blue actinic light. Results represent the mean of 3 biological replicates; error bars represent standard deviation, and significance was determined using two-way ANOVA. **D.** Clear-native PAGE (CN-PAGE) on thylakoid membrane extracts from cells grown under 70 μmol photons m^−2^ s^−1^. Extraction was done on mid-log phase (OD730=0.6) cells. Thylakoid membranes were quantified according to the chlorophyll content. Samples containing the same amount of chlorophyll were loaded on the gel. (1) Wild-type (2) ΔAdo (3) ΔAdo-FAP Control (4) ΔAdo-FAP.

We then investigated the effects of alka(e)ne deficiency on photosynthesis. For this purpose, we performed chlorophyll fluorescence measurements to determine the max PSII efficiency (Fv/Fm), the operating PSII quantum yield under actinic light (ΦPSII), and energy dissipation by non-photochemical quenching (NPQ). O_2_ measurements were performed simultaneously by using membrane inlet mass spectrometry (MIMS) in the presence of ^18^O-labeled O_2_ to determine gross O_2_ production rates by PSII and O_2_ consumption rates in the light (28). These measurements were performed in response to different illumination regimes, with cells grown under either low light (10 μmol photons m^-2^ s^-1^) or relatively high light (70 μmol photons m^-2^ s^-1^). When the different *Synechocystis* strains were grown under low light, no significant changes in Fv/Fm, ΦPSII, NPQ and O_2_ exchange were observed among all the strains (Fig. 2B-C, Supplementary Figure 2). When cultivated under higher light intensity, the alka(e)ne-deficient strains exhibited notably lower Fv/Fm and ΦPSII values, along with increased NPQ, compared to the WT under actinic illumination. This effect was reversed in the functionally complemented strain expressing FAP (Fig. 2B-C). In line with chlorophyll fluorescence measurements, both gross and net O_2_ production rates were decreased in the alka(e)ne-deficient strains grown under high light intensity (Fig. 2C), while O_2_ uptake rate being mostly unaffected (Supplementary Figure 2). We then analyzed the distribution of chlorophyll protein complexes by performing a native PAGE on thylakoids membrane prepared from the cells cultivated under high light. Whereas PSII-complexes were mostly unaffected, we observed a dramatic reduction in the PSI trimer content in alka(e)nes-deficient strains (Fig. 2D), thus indicating that PSI trimerization and/or stabilization might depend on the presence of alka(e)nes in the thylakoid membrane. We conclude from these results that the absence of alka(e)nes strongly affects the photosynthetic machinery and its functioning under high light.

### The cyanobacterial lipidome is drastically remodeled in the absence of alka(e)nes

Given the close link between alka(e)ne and fatty acid biosynthetic pathways (Supplementary Figure 3), and the altered ultrastructures of photosynthetic membranes observed in alka(e)ne-deficient strains, we sought to analyze in depth the effect of alka(e)ne-deficiency on the lipidome. We first analyzed changes in the fatty acid content of the various strains cultivated under either low light (10 μmol photons m^-2^ s^-1^) or high light (70 μmol photons m^-2^ s^-1^). Under low light, alka(e)ne-deficient strains exhibited higher contents of C16:0, C16:1, and C18:1 fatty acids as compared to WT and complemented strains (Fig. 3A). In contrast, under high light, alka(e)ne-deficient strains showed lower contents of C16:0 and C16:1 than the WT and complemented strains. However, the proportion of saturated, unsaturated and cyclo fatty acids remained unchanged among the strains under both low and high light (Supplementary Figure 4).

**Fig. 3.**
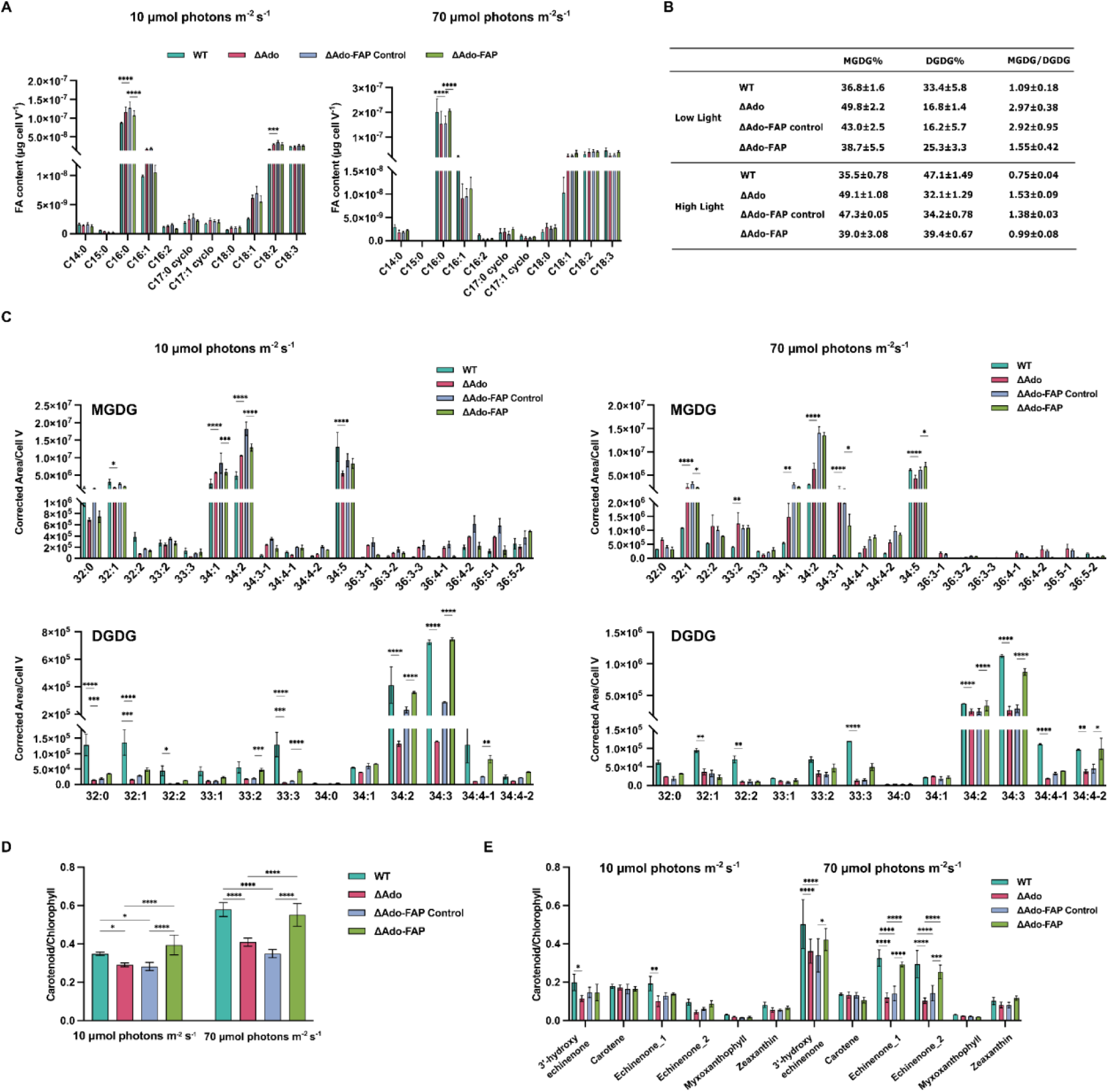
Fatty acid content, composition in major membrane glycerolipids, and carotenoid to chlorophyll ratio under low and high light intensities. **A.** Fatty acid content in *Synechocystis* strains. The contents of different fatty acid species were quantified by cell transmethylation and GC-MS and normalized to cell volume representing biomass. **B.** The proportion of MGDG and DGDG in total lipid content, and MGDG to DGDG ratio in *Synechocystis* strains. Absolute quantification of each lipid class was performed using HPTLC and densitometry. **C.** Lipidomics analysis on DGDG and MGDG species in different strains. Results represent the mean of 3 biological replicates; error bars represent standard deviation. **D.** Spectrophotometer measured carotenoids to chlorophyll content ratio in low light and high light cultivated strains. Pigment extraction was done using 90% pre-cold methanol. **E.** Different Carotenoid species to chlorophyll ratio measured by LC-MS. Left panel shows pigment ratio under low light and right panel shows pigment ratio under high light. Carotenoids and chlorophyll were extracted from mid-log phase (OD_730_=0.6) culture using 90% pre-cold methanol, then the extracts were measured on LC-MS. All results represent the mean of 3 biological replicates; error bars represent standard deviation, and significance was determined using ANOVA.

The composition of major membrane lipid classes was then analyzed using high-performance thin-layer chromatography (HPTLC). Alka(e)ne-deficient strains consistently exhibited higher contents of the non-bilayer-forming lipid monogalactosyldiacylglycerol (MGDG) and lower contents of the bilayer-forming lipid digalactosyldiacylglycerol (DGDG) than the WT and complemented strains, irrespective of light conditions, while sulfoquinovosyldiacylglycerol (SQDG) and phosphatidylglycerol (PG) contents remained relatively constant (Fig. 3B, Supplementary Figure 5). To gain further insights into the dynamic changes of MGDG and DGDG lipid species, we performed a comprehensive LC-MS lipidomic analysis of glycerolipid molecular species which provides a relative quantification and allows to compare the dynamics of specific lipid species across different strains or experimental conditions (Fig. 3C). This analysis showed that the content of all DGDG species analyzed decreased in alka(e)ne-deficient strains regardless of light conditions (Fig. 3C). In parallel, the amount of most MGDG species increased in the absence of alka(e)nes. Notably, the expression of FAP led to a partial recovery of several DGDG species (*e.g.* 32:1 and 33:3), with a nearly complete recovery observed for some of them (*e.g.* 34:2 and 34:3). For MGDG species, FAP expression facilitated the recovery of nearly half of these lipid species, with no observable recovery for others (*e.g*. 32:1 and 34:2).

The major changes observed in the lipid composition of alka(e)ne-deficient strains prompted us to analyze the potential impact on pigment composition, since carotenoids are primarily located within the thylakoid membranes, where they are integrated alongside chlorophyll molecules and other pigments in the photosynthetic complexes. Carotenoids participate in light absorption and photoprotection (29), and have been shown to respond to challenging environmental conditions, such as intense light and high temperatures (30, 31). In low light conditions, the carotenoid content of alka(e)ne-deficient strains was slightly lower than the WT strain (Fig. 3D), mainly due to the decrease in 3-hydroxy echinenone and two echinenone isoforms (Fig. 3E). Under high light conditions, the carotenoid content of the WT strain increased (Fig. 3D), due to the increase of 3-hydroxy echinenone and of the two echinenone isoforms (Fig. 3E), the increase being reduced in alka(e)ne-deficient strains. Complemented strains expressing FAP showed recovery of all carotenoids (Fig. 3E) under both light conditions. We did not observe a significant change in chlorophyll content from cultures grown under the same light condition (Supplementary figure 6), indicating that observed changes in carotenoid do not reflect a decline in all photosynthetic pigments.

## Discussion

In this study, we investigated the physiological role of alka(e)nes in relation to the photosynthetic function, using the cyanobacterium *Synechocystis* PCC6803 as a model organism. The absence of alka(e)nes provoked remarkable structural and physiological changes, including an altered pigment content, a profound remodeling of the glycerolipid composition of the thylakoid membranes, and a growth defect under high light intensity. Specifically, when alka(e)ne deficient strains were cultivated under high light, the photosynthetic activity was reduced, and the non-photochemical quenching increased. The lipidomics analysis revealed a decrease in DGDG and an increase in MGDG levels together with a decrease in the carotenoid content in alka(e)ne-deficient strains in both low and high light conditions. All of the observed phenotypes could be either partially or completely restored when the alka(e)ne-producing photoenzyme FAP was expressed in the mutant strains.

### Alka(e)nes deficiency affects thylakoid membrane dynamics

A molecular dynamics simulation has suggested that the alka(e)nes located in the center of bilayer membrane would contribute to membrane flexibility thus facilitating membrane curvature (24). Therefore, the decreased MGDG to DGDG ratio in the absence of alka(e)ne regardless of cultivation light suggests a primary compensatory effect that may stem from the lack of membrane flexibility and curvature. MGDG is one of the most abundant lipids in thylakoid membranes and has a cone-shaped molecular structure. This shape tends to promote the formation of non-bilayer (hexagonal II) phases, which are more fluid. MGDG is known to play a crucial role in maintaining the fluidity of the thylakoid membrane and is vital for the proper functioning of the photosynthetic machinery (32, 33). On the other hand, DGDG, with its more cylindrical shape due to the addition of another galactose group, favors bilayer formation. DGDG can contribute to membrane rigidity and stability, especially under stress conditions. For instance, under phosphate-limited conditions, DGDG can substitute for phospholipids in various membranes to ensure membrane stability (34, 35). A study employing neutron diffraction on thylakoid extracts revealed the important role of DGDG in facilitating membrane stacking (36). The diminished DGDG content observed in alka(e)ne-deficient strains would then lead to thylakoids being more dispersed rather than stacked, creating the illusion of an increased thylakoid presence.

Absence of alka(e)nes did not only impact the glycerolipid composition in the thylakoid membranes, it also had an effect on the associated carotenoids. Carotenoids are tetra-terpenoid molecules found in all photosynthetic organisms, playing a key role in enhanced light absorption and energy dissipation during photosynthesis (37). In our alka(e)ne-deficient strains, lower carotenoid content was observed regardless of light and CO2 availability (Supplementary figure 7). As for the MGDG/DGDG ratio, the alteration of the carotenoid content could be a compensatory effect directly from the change of membrane fluidity, or result from changes of the lipid environment. It has been shown that in *Arabidopsis thaliana*, although most of the carotenoids are associated with light-harvesting complexes, around 15% of the carotenoids are still freely distributed throughout the thylakoid membrane (38, 39), and the lutein/carotene ratio significantly affects thylakoid membrane fluidity (40). In addition, the change in the carotenoid content observed in alka(e)nes-deficient mutants may explain the observed effect on the S-layer (Fig. 1C). Indeed, the absence of an S-layer in a xanthophyll-deficient *Synechocystis* mutant (41) was explained by the fact that xanthophylls can provide a proper environment in the outer membrane for anchoring S-layer proteins to lipopolysaccharides.

### Alka(e)nes indirectly affect photosynthesis resilience

In addition to all the changes in the structure and lipid composition of the thylakoid membranes, we observed reduced photosynthetic properties when alka(e)ne-deficient strains were exposed to high light. The higher light sensitivity of photosynthesis is likely an indirect consequence of the altered membrane lipid composition resulting from alkane(e) deficiency. Indeed, previous studies in cyanobacteria and plants have shown that changes in DGDG content affect photosynthesis functionality (42, 43). *Synechocystis* cells lacking DGDG experience stunted growth under high light conditions, potentially impacting dynamic processes in photosynthesis such as the PSII repair process (44). Alternatively, the observed changes in thylakoid stacking in alka(e)ne-deficient strains might induce an increased susceptibility to light stress. Enhanced stacking of thylakoids can decrease the effective surface area exposed to high light intensities, serving as a defense against photodamage (45, 46). Furthermore, the increased high light sensitivity of alka(e)ene deficient strains may also partly result from changes in the carotenoid composition. It has been shown that carotenoid content, specifically echinenone, increased sharply under high light illumination in both algae and cyanobacteria (47, 48). The sharp increase of echinenone observed in *Synechocystis* WT in response to high light (Fig. 3D) was strongly diminished in the alka(e)ne-deficient strains (Fig. 3D), which could lead to a higher light-sensitivity of PSII. Indeed, it has been reported that cells lacking zeaxanthin, echinenone, and myxoxanthophyll show a significant decrease in the capacity for repair of PSII under strong light (48).

The decrease in the amount of PSI trimers observed in alka(e)ne-deficient strains likely results from changes in the lipid and carotenoid contents. Indeed, it has been shown in *Arabidopsis thaliana* that DGDG deficiency resulted in a restructuring of the thylakoid domain, causing significant changes in the PSI structure (49). This structural shift hinders intersystem electron transport and disrupts the regulation of energy distribution via state transitions. The importance of carotenoids in photosynthetic organisms was further revealed with recent findings highlighting their crucial role in the assembly of photosynthetic complexes (50). For instance, xanthophylls, such as echinenone and zeaxanthin, have been pinpointed as instrumental in the fine-tuning of PSI trimerization (51). This is further illuminated by the crystal structure of the pigment-protein PSI trimeric complex resolved to 2.5 Å using X-ray crystallography (52). Here, echinenone is predominantly situated at the interface between two monomers, with zeaxanthin positioned on the exterior of each monomer, and a single zeaxanthin per monomer. This configuration provides insight into our observations: strains with diminished xanthophyll content exhibit fewer PSI trimers, which is in line with a previous study where the *Synechocystis* mutant lacking xanthophylls showed fewer PSI trimers (41).

A previous study showed the absence of alka(e)nes results in minor effects on photosynthetic performance of *Synechocystis* cells (24). In this work the authors used an intermediate light intensity (30 μmol photons m^-2^ s^-1^) as compared to our own conditions (10 and 70 μmol photons m^-2^ s^-1^ for low light and high light respectively). Moreover, the authors did supplement the culture medium with bicarbonate (24), whereas no additional carbon was supplied in our experiments. It seems likely that both light and carbon supply affect the high light phenotype of alka(e)ne deficient strains; indeed, we observed that when grown in air supplemented with 1% CO2 the photosynthetic capacity of alka(e)ne deficient strains is not affected even under high light (70 μmol photons m^-2^ s^-1^) condition (Supplementary figure 8).

Based on the observed phenotypes, we propose the following scenario (Fig. 4): the absence of alka(e)nes would induce a change in membrane physical properties, such as fluidity, which would trigger compensatory mechanisms including a decrease in DGDG and an increase in MGDG and a change in the carotenoid content. This would in turn induce an instability of PSI trimers and a disturbance of the PSII repair, thus affecting the functioning of the electron transport chain and compromising growth under high light.

**Fig. 4.**
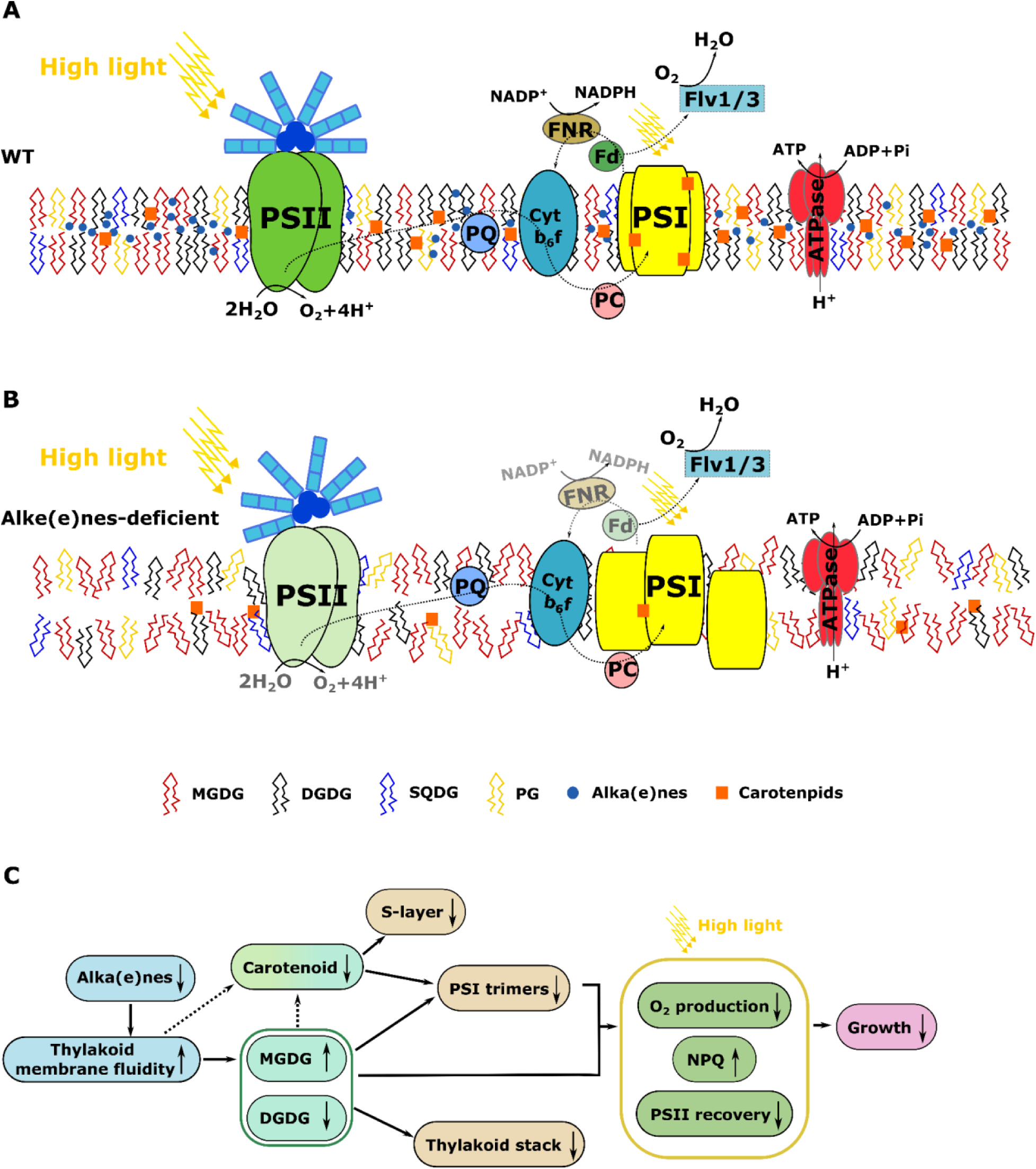
Scheme of the proposed scenario due to alka(e)nes deficiency. **A.** Thylakoid membrane arrangement in wild-type *Synechocystis* cells. **B.** Thylakoid membrane arrangement in alka(e)nes-deficient *Synechocystis* cells. PQ: Plastoquinone, Cytb6f: Cytochrome b6f complex, FNR: Ferredoxin—NADP (+) reductase, Fd: Ferredoxins, PC: plastocyanin, Flv1/3: Flavodiiron proteins Flv1 and Flv3. **C**. Proposed chain of effects resulting from alka(e)nes deficiency.

In conclusion, the current work sheds light on the ecological significance of alka(e)nes, which play a part in membrane lipid homeostasis and are essential for the resilience of photosynthesis under high light. Uncovering the physiological role of alka(e)nes provides insights to a better understanding of the widespread presence of genes encoding alka(e)nes-synthesizing enzymes in the genomes of cyanobacteria and microalgae, organisms of major ecological importance in the global capture of CO_2_ by aquatic ecosystems.

## Materials and Methods

### Strains and cultivation conditions

*Escherichia coli* strains DH5α and XL1-Blue were used for plasmids construction. The cells were grown at 37°C in LB medium supplemented with corresponding antibiotics for selection. Glucose-tolerant *Synechocystis* PCC6803 was used as background strain in this study, and BG11 medium used here was always buffered to pH=7.8 by 20 mM TES. To generate knock-out mutants, the target gene/operon was replaced with an antibiotic resistance gene through homologous recombination. This was facilitated by two homologous overhangs at both the 5’ and 3’ ends of the target operon or gene, with each overhang being 1000 bp in length. To express FAP in the *slr1993* loci, the *slr1993* gene was replaced by FAP and a spectinomycin resistance gene using homologous recombination. Genetic modifications of all engineered *Synechocystis* strains are explained in Supplementary table 1. Natural transformation was performed using mid-log phase *Synechocystis* culture. For each engineered strain, 3 independent biological replicates were used. Culture maintenance was done using BG11 liquid media or agar plates with corresponding antibiotics. In order to remove the stresses from different antibiotics, BG11 medium without any antibiotics was used to perform all the experimental culturing. Genotype stability from cultures without antibiotics were confirmed from time to time. Liquid culture cultivation was done in E-flasks with a light source from fluorescent lamp Fluora Osram, and three different light intensities were used: 10 μmol photons m^-2^ s^-1^, 30 μmol photons m^-2^ s^-1^, and 70 μmol photons m^-2^ s^-1^. Cell growth test on BG11-agar plates was performed under three light intensities: 20 μmol photons m^-2^ s^-1^, 50 μmol photons m^-2^ s^-1^, and 200 μmol photons m^-2^ s^-1^. Cell size, number, and volume were measured at each sampling time using a Multisizer Beckman Coulter.

### Chlorophyll α, and carotenoid measurements

The overall concentration determination of chlorophyll a and carotenoid was conducted on the 90% methanol extracts obtained from 1 mL of each *Synechocystis* culture. Absorbance for chlorophyll concentration was taken at 665 nm, while for carotenoid concentration, it was taken at 470 nm. Additionally, absorbance at 720 nm was measured to account for background. The calculations were based on the following equations: Chla[µg/ml] = 12.94*(A665-A720) and Carotenoids[µg/ml] = [1000*(A470-A720)-2.86*(Chla/221)]. For the assessment of chlorophyll and individual carotenoid species content, pigment extracts from four milliliters of each *Synechocystis* culture was utilized on LC-MS.

### Alka(e)nes and fatty acids analysis

To determine the content of alka(e)nes and fatty acids, 4 mL of each *Synechocystis* culture in mid-log-phase was sampled. It was then centrifuged in a glass tube at 3000 rpm for 10 min. Following this, direct transmethylation was carried out on the harvested cell pellet. Into this pellet, 1 mL of 5% (v/v) H_2_SO_4_ in methanol, 10 µg of internal standards (*n*-hexadecane and *n*-heptadecanoic acid), and 10 µL of 10% (w/v) butylated hydroxytoluene in methanol (BHT,an antioxidant) were added. The resulting mixture in the glass tube was homogenized through sonication. The sealed tube was then heated at 85°C for 1.5 hours. Once it had cooled to room temperature, 1.5 mL of 0.9% (w/v) NaCl was introduced to assist in biphasic separation. Additionally, 500 µL of hexane was added to extract the fatty acid methyl esters and the alka(e)nes. A subsequent centrifugation at 4000 rpm for 4 min was performed, and the hexane layer was collected for further analysis. The heptane phase was injected in an Agilent 7890A gas chromatograph coupled to a Flame Ionization Detector (FID) and an Agilent 5975C mass spectrometer. The column was a Zebron 7HG-G007-11 (Phenomenex) polar capillary column (30 m length, internal diameter of 0.25 mm, film thickness 0.25 mm). Dihydrogen was the carrier gas, with a flow rate maintained at 1 mL min^-1^. Fatty acids and alka(e)nes were identified based on their retention time and mass spectrum and quantified based on their FID peak area and internal standards.

### Lipid extractions

An isopropanol/MTBE method was applied to extract lipids for both LC-MS and HPTLC analysis. Briefly, cultures of the same total cell volume were subjected to centrifugation in a glass tube at 3000 g and 4°C for 4 min, and the supernatant was subsequently discarded. For each sample, 1 mL of pre-heated isopropanol, supplemented with 10 µL of 1% (w:v) BHT and 1 µL formic acid, was added to the glass tube. Resuspension of the cells was achieved through vigorous vortexing and sonication. For the LC-MS samples, an internal standard of 10 µL was incorporated, while the HPTLC samples did not require an internal standard. The glass tubes, sealed with screw caps, were heated at 85 °C for 10 min to ensure lipase inactivation. Once the samples reached room temperature, 3 mL of methyl-tert-butyl ether (MTBE) was added to each tube and they were shaken vigorously for 2 minutes. A mixture of 1 mL of 0.9% NaCl and 1 µL formic acid was subsequently added to each sample, followed by vigorous shaking. Phase separation was achieved by centrifuging the samples at 3500 g and 4°C for 2 min. The upper phase was carefully transferred to a new glass tube. A second extraction was initiated by adding 1 mL of MTBE to the remaining lower phase. The upper phases from these two procedures were combined, and the solvent was allowed to evaporate under a stream of nitrogen gas. For LC-MS analysis, lipids were redissolved in 500 µL of a solvent mixture acetonitrile/isopropanol/ammonium formate (65:30:5, v/v/v). For HPTLC, lipids were resuspended in 400 µL of a freshly prepared chloroform:methanol mixture 2:1 (v:v).

### Lipidomics analysis by LC-MS

Lipid molecular species were analyzed on an ultimate RS 3000 UPLC system (Thermo Fisher, Waltham, MA, USA) connected to a quadrupole-time-of-flight (QTOF) 5600 mass spectrometer (AB Sciex, Framingham, MA, USA) equipped with an ion source operating in negative mode. Lipid extracts were first separated on a KinetexTM (Kinetex, Atlanta, GA, USA) C18 2.1 × 150 mm 1.7 μm column (Phenomenex, Torrance, CA, USA). Two solvent mixtures, acetonitrile−water (60:40, v/v) and isopropanol−acetonitrile (90:10, v/v), both containing 10 mM ammonium formate at pH 3.8, were used as eluent A and B respectively. The elution was performed with a gradient of 32 min; eluant B was increased from 27 to 97% in 20 min then maintained for 5 min, solvent B was decreased to 27% and then maintained for another 7 min for column re-equilibration. The flow-rate was 0.3 mL.min-1 and the column oven temperature was maintained at 45°C. Lipid identification was based on retention time and on mass accuracy peaks from the MS survey scan compared with theoretical masses and fragment ions from MS/MS scan. For each molecular species, relative quantification was achieved with Multiquant software (AB Sciex) on the basis of the intensity of the molecular ion of one adduct.

### Analysis of lipid classes by HPTLC

Quantification of polar lipids content on HPTLC was carried out as follows: Lipid extracts and polar lipids standards at varying concentrations were applied onto silica gel 60 F254 HPTLC plates (Merck KGaA, Germany) using an ATS 5 automatic TLC sampler (Camag, Switzerland). Subsequently, these plates were developed in an ADC2 automatic developing chamber (Camag) with a solvent mixture of acetone:toluene:water in a ratio of 19:30:8 (v:v:v). Once developed, the plate was allowed to dry thoroughly under a hood. It was then dipped in a CuSO4 reagent for 6 seconds. After this treatment, the plate was subjected to heating at 141°C for 30 minutes. Scanning of the plate was performed using a TLC Scanner 3 operated with WinCATs software (Camag). Lipid classes were quantified based on a standard range for each class.

### Chlorophyll fluorescence and oxygen evolution measurements

Chlorophyll fluorescence and oxygen evolution were simultaneously measured using a pulsed amplitude modulation (PAM) fluorimeter (Dual-PAM 100) coupled with the membrane inlet mass spectrometry (MIMS) setup (28). Two mL of cell suspension at a final chlorophyll content of 5 μg Chl ml^-1^ were placed in a flat bottom stirred temperature controlled (30°C) cuvette placed in darkness for 10 min, and 10 µL of 0.5 M NaHCO_3_ were added. The cell suspension was bubbled with ^18^O-enriched O_2_ (with 99% ^18^O_2_ isotope content from Euriso-Top) until concentrations of ^16^O_2_ and ^18^O_2_ reached approximately equal values. Both oxygen exchange and chlorophyll fluorescence were then measured in the dark for 3 min and then in response to stepwise increase of blue actinic light intensity up to 6200 μmol photons m^-2^ s^-1^. NPQ was determined as (Fm-Fm’)/Fm’ based on the chlorophyll fluorescence Fm measured in darkness. Meanwhile, O_2_ uptake, gross O2 production and net O2 evolution rates were measured on the same sample using MIMS as described in a previous study (28).

### Thylakoid extraction and native PAGE

Cells with an OD_730_ ranging from 0.5 to 0.8, were harvested from a fifty-milliliter sample via centrifugation at 2300 g and 4°C for 10 minutes. The resultant pellet was then resuspended in 150 µL of thylakoid buffer (containing 25 mM MES-NaOH at pH 6.5, 10 mM CaCl_2_, 10 mM MgCl_2_, and 25% (w/v) glycerol) and relocated to a 2 mL screw cap tube. A set of glass beads from a PCR-tube was introduced to the tube, followed by cell disruption using a liquid nitrogen-paired BeadBeater. Subsequently, 200 µL of the thylakoid buffer was added, the mixture was vortexed twice, and the supernatant was transferred to a new, chilled Eppendorf tube. This procedure was repeated five times. To separate any remaining unbroken cells and glass beads, the mixture was centrifuged at 4000 g for a minute. The clear supernatant was then moved to another cooled Eppendorf tube and subjected to further centrifugation at 16000 g and 4°C for 20 minutes. After discarding the supernatant, the pellet was resuspended in 200 µL of thylakoid buffer with the aid of a stainless-steel piston and vortexing. For membrane solubilization, a tenth of the volume of freshly prepared 10% (v/v) n-Dodecyl-β-D-maltoside (DM) was added to the samples, which were then incubated on ice for 5 minutes. Chlorophyll extraction from the samples was performed using methanol, and its concentration was determined spectrophotometrically. Afterward, the solubilized membrane sample, containing 10 µg of chlorophyll, was centrifuged again under the same conditions. The resultant supernatant was transferred to a new chilled Eppendorf tube and mixed with 10% (v/v) deoxycholate (DOC) to achieve a final DOC concentration of 0.5%. This mixture was then loaded onto a Mini-PROTEAN TGX Gel (BioRad) for clear native PAGE. The buffer system used for the native PAGE consisted of an upper buffer (with 0.05 M tricine, 15 mM Bis-Tris/HCl at pH 7, 0.05% (v/v) DOC, and 0.02% (v/v) DM) and a lower buffer (0.05 M Bis-Tris/HCl at pH 7). Importantly, all extraction and native PAGE procedures were conducted at 4°C and shielded from light.

### Transmission electron microscopy

*Synechocystis* cells in the mid-log phase (OD_730_=0.5-0.8) were subjected to centrifugation at 3000 g and 4°C for 10 minutes. Subsequently, the cells were fixed in 2.5% (v/v) glutaraldehyde combined with 1% (v/v) paraformaldehyde and 0.1 M PIPES. This fixation process was carried out for 60 minutes on a shaker at room temperature. For embedding, the cells underwent a series of treatments: 1) Rinsing in 0.1 M PB for 10 minutes. 2) A 120-minute incubation in 0.1 M PB containing 1% (v/v) Osmium tetroxide was applied. 3) Another 10-minute rinse in 0.1 M PB followed., 4) The cells were sequentially exposed to increasing concentrations of ethanol (ranging from 50% to 100% (v/v), with each concentration applied for 10 minutes. 5) A 30-minute incubation in a 100% ethanol:LR white resin mixture in a 1:1 (v/v) ratio was done. 6) An overnight immersion in LR white resin took place, after which the medium was replaced with fresh LR white acrylic resins. 7) Curing at 60 °C for 48 hours. Between each of the aforementioned steps, the cells were settled by centrifuging at 2300 g for 2 minutes. Ultra-thin sections, between 70-80 nm in thickness, were sliced using a Leica UCT Ultramicrotome and carefully positioned on grids. These grids were then treated with stains: 2% (v/v) Uranyl Acetate followed by 3% (v/v) Reynolds lead citrate. Finally, the stained specimens were analyzed using a TEM (FEI Tecnai G2) operating at 80kV.

## Acknowledgements

This work was funded by the ANR project Photoalkane (ANR-18-CE43-0008). We thank the Torsten Thuréns stiftelse för miljövänlig forskning for providing financial support for the transmission electron microscopy experiment. We also thank the Federation of European Microbiological Societies (FEMS) for providing a research training grant for completing part of the photosynthetic capacity measurements. Excellent internship work of Ms. Emma Calikanzaros who contributed to part of this study is also acknowledged. We also acknowledge the European Union Regional Developing Fund (ERDF), the Région Provence Alpes Côte d’Azur, the French Ministry of Research and the CEA for funding the HelioBiotec platform.

## References

1. P. Laurent, J.-C. Braekman, D. Daloze, J. Pasteels, Biosynthesis of defensive compounds from beetles and ants. European J. Org. Chem. 2003, 2733–2743 (2003).

2. M. A. Major, G. J. Blomquist, Biosynthesis of hydrocarbons in insects: Decarboxylation of long chain acids ton-alkanes in *Periplaneta*. Lipids 13, 323–328 (1978).

3. G. J. Blomquist, M. D. Ginzel, Chemical Ecology, Biochemistry, and Molecular Biology of Insect Hydrocarbons. Annu. Rev. Entomol. 66, 45–60 (2021).

4. H. Chung, S. B. Carroll, Wax, sex and the origin of species: Dual roles of insect cuticular hydrocarbons in adaptation and mating. Bioessays 37, 822–830 (2015).

5. T. H. Yeats, J. K. C. Rose, The formation and function of plant cuticles. Plant Physiol. 163, 5–20 (2013).

6. S. B. Lee, M. C. Suh, Regulatory mechanisms underlying cuticular wax biosynthesis. J. Exp. Bot. 73, 2799–2816 (2022).

7. B. Valderrama, “Chapter 13 Bacterial hydrocarbon biosynthesis revisited” in Studies in Surface Science and Catalysis, R. Vazquez-Duhalt, R. Quintero-Ramirez, Eds. (Elsevier, 2004), pp. 373–384.

8. N. A. Herman, W. Zhang, Enzymes for fatty acid-based hydrocarbon biosynthesis. Curr. Opin. Chem. Biol. 35, 22–28 (2016).

9. D. Sorigué, et al., Microalgae Synthesize Hydrocarbons from Long-Chain Fatty Acids via a Light-Dependent Pathway. Plant Physiol. 171, 2393–2405 (2016).

10. D. Sorigué, et al., An algal photoenzyme converts fatty acids to hydrocarbons. Science 357, 903–907 (2017).

11. P. P. Samire, et al., Autocatalytic effect boosts the production of medium-chain hydrocarbons by fatty acid photodecarboxylase. Sci Adv 9, eadg3881 (2023).

12. D. Sorigué, et al., Mechanism and dynamics of fatty acid photodecarboxylase. Science 372 (2021).

13. S. L. Y. Moulin, et al., Fatty acid photodecarboxylase is an ancient photoenzyme that forms hydrocarbons in the thylakoids of algae. Plant Physiol. 186, 1455–1472 (2021).

14. W. Wang, X. Liu, X. Lu, Engineering cyanobacteria to improve photosynthetic production of alka(e)nes. Biotechnol. Biofuels 6, 69 (2013).

15. M. Xie, W. Wang, W. Zhang, L. Chen, X. Lu, Versatility of hydrocarbon production in cyanobacteria. Appl. Microbiol. Biotechnol. 101, 905–919 (2017).

16. Y. Hayashi, M. Arai, Recent advances in the improvement of cyanobacterial enzymes for bioalkane production. Microb. Cell Fact. 21, 256 (2022).

17. I. S. Yunus, et al., Synthetic metabolic pathways for conversion of CO2 into secreted short-to medium-chain hydrocarbons using cyanobacteria. Metab. Eng. 72, 14–23 (2022).

18. A. Schirmer, M. A. Rude, X. Li, E. Popova, S. B. del Cardayre, Microbial biosynthesis of alkanes. Science 329, 559–562 (2010).

19. D. Mendez-Perez, M. B. Begemann, B. F. Pfleger, Modular synthase-encoding gene involved in α-olefin biosynthesis in *Synechococcus* sp. strain PCC 7002. Appl. Environ. Microbiol. 77, 4264–4267 (2011).

20. R. C. Coates, et al., Characterization of cyanobacterial hydrocarbon composition and distribution of biosynthetic pathways. PLoS One 9, e85140 (2014).

21. D. J. Lea-Smith, et al., Contribution of cyanobacterial alkane production to the ocean hydrocarbon cycle. Proc. Natl. Acad. Sci. U. S. A. 112, 13591–13596 (2015).

22. C. R. Love, et al., Microbial production and consumption of hydrocarbons in the global ocean. Nat Microbiol 6, 489–498 (2021).

23. S. Klähn, et al., Alkane Biosynthesis Genes in Cyanobacteria and Their Transcriptional Organization. Front Bioeng Biotechnol 2, 24 (2014).

24. D. J. Lea-Smith, et al., Hydrocarbons Are Essential for Optimal Cell Size, Division, and Growth of Cyanobacteria. Plant Physiol. 172, 1928–1940 (2016).

25. B. M. Berla, R. Saha, C. D. Maranas, H. B. Pakrasi, Cyanobacterial Alkanes Modulate Photosynthetic Cyclic Electron Flow to Assist Growth under Cold Stress. Sci. Rep. 5, 14894 (2015).

26. T. Yamamori, H. Kageyama, Y. Tanaka, T. Takabe, Requirement of alkanes for salt tolerance of Cyanobacteria: characterization of alkane synthesis genes from salt-sensitive *Synechococcus elongatus* PCC7942 and salt-tolerant Aphanothece halophytica. Lett. Appl. Microbiol. 67, 299–305 (2018).

27. N. Li, et al., Conversion of fatty aldehydes to alka(e)nes and formate by a cyanobacterial aldehyde decarbonylase: cryptic redox by an unusual dimetal oxygenase. J. Am. Chem. Soc. 133, 6158–6161 (2011).

28. A. Burlacot, F. Burlacot, Y. Li-Beisson, G. Peltier, Membrane Inlet Mass Spectrometry: A Powerful Tool for Algal Research. Front. Plant Sci. 11, 1302 (2020).

29. D. Kirilovsky, C. A. Kerfeld, The orange carotenoid protein in photoprotection of photosystem II in cyanobacteria. Biochim. Biophys. Acta 1817, 158–166 (2012).

30. M. Havaux, Carotenoids as membrane stabilizers in chloroplasts. Trends Plant Sci. 3, 147–151 (1998).

31. N. Dhami, J. E. Drake, M. G. Tjoelker, D. T. Tissue, C. I. Cazzonelli, An extreme heatwave enhanced the xanthophyll de-epoxidation state in leaves of Eucalyptus trees grown in the field. Physiol. Mol. Biol. Plants 26, 211–218 (2020).

32. D. Marsh, Components of the lateral pressure in lipid bilayers deduced from HII phase dimensions. Biochim. Biophys. Acta 1279, 119–123 (1996).

33. J. Hoyo, E. Guaus, J. Torrent-Burgués, Monogalactosyldiacylglycerol and digalactosyldiacylglycerol role, physical states, applications and biomimetic monolayer films. Eur. Phys. J. E Soft Matter 39, 39 (2016).

34. H. Härtel, P. Dormann, C. Benning, DGD1-independent biosynthesis of extraplastidic galactolipids after phosphate deprivation in *Arabidopsis*. Proc. Natl. Acad. Sci. U. S. A. 97, 10649–10654 (2000).

35. M. X. Andersson, P. Dörmann, “Chloroplast Membrane Lipid Biosynthesis and Transport” in *Plant Cell Monographs*, Plant cell monographs., (Springer Berlin Heidelberg, 2008) 10.1007/7089_2008_18.

36. B. Demé, C. Cataye, M. A. Block, E. Maréchal, J. Jouhet, Contribution of galactoglycerolipids to the 3-dimensional architecture of thylakoids. FASEB J. 28, 3373–3383 (2014).

37. F. Pagels, V. Vasconcelos, A. C. Guedes, Carotenoids from Cyanobacteria: Biotechnological Potential and Optimization Strategies. Biomolecules 11 (2021).

38. L. Dall’Osto, S. Cazzaniga, M. Havaux, R. Bassi, Enhanced photoprotection by protein-bound vs free xanthophyll pools: a comparative analysis of chlorophyll b and xanthophyll biosynthesis mutants. Mol. Plant 3, 576–593 (2010).

39. T. Zakar, H. Laczko-Dobos, T. N. Toth, Z. Gombos, Carotenoids Assist in Cyanobacterial Photosystem II Assembly and Function. Front. Plant Sci. 7, 295 (2016).

40. M. Bykowski, et al., Too rigid to fold: Carotenoid-dependent decrease in thylakoid fluidity hampers the formation of chloroplast grana. Plant Physiol. 185, 210–227 (2021).

41. T. Zakar, et al., Lipid and carotenoid cooperation-driven adaptation to light and temperature stress in *Synechocystis* sp. PCC6803. Biochim. Biophys. Acta Bioenerg. 1858, 337–350 (2017).

42. H. Härtel, H. Lokstein, P. Dörmann, B. Grimm, C. Benning, Changes in the composition of the photosynthetic apparatus in the galactolipid-deficient dgd1 mutant of Arabidopsis thaliana. Plant Physiol. 115, 1175–1184 (1997).

43. I. Sakurai, N. Mizusawa, H. Wada, N. Sato, Digalactosyldiacylglycerol is required for stabilization of the oxygen-evolving complex in photosystem II. Plant Physiol. 145, 1361–1370 (2007).

44. N. Mizusawa, I. Sakurai, N. Sato, H. Wada, Lack of digalactosyldiacylglycerol increases the sensitivity of *Synechocystis* sp. PCC 6803 to high light stress. FEBS Lett. 583, 718–722 (2009).

45. J. M. Anderson, E. M. Aro, Grana stacking and protection of Photosystem II in thylakoid membranes of higher plant leaves under sustained high irradiance: An hypothesis. Photosynth. Res. 41, 315–326 (1994).

46. W. H. J. Wood, et al., Dynamic thylakoid stacking regulates the balance between linear and cyclic photosynthetic electron transfer. Nat Plants 4, 116–127 (2018).

47. A. Pajot, et al., Light-response in two clonal strains of the haptophyte Tisochrysis lutea: Evidence for different photoprotection strategies. Algal Research 69, 102915 (2023).

48. T. Izuhara, I. Kaihatsu, H. Jimbo, S. Takaichi, Y. Nishiyama, Elevated Levels of Specific Carotenoids During Acclimation to Strong Light Protect the Repair of Photosystem II in *Synechocystis* sp. PCC 6803. Front. Plant Sci. 11, 1030 (2020).

49. A. G. Ivanov, et al., Digalactosyl-diacylglycerol deficiency impairs the capacity for photosynthetic intersystem electron transport and state transitions in *Arabidopsis thaliana* due to photosystem I acceptor-side limitations. Plant Cell Physiol. 47, 1146–1157 (2006).

50. T. N. Tóth, et al., Carotenoids are essential for the assembly of cyanobacterial photosynthetic complexes. Biochim. Biophys. Acta 1847, 1153–1165 (2015).

51. S. Vajravel, et al., Zeaxanthin and echinenone modify the structure of photosystem I trimer in Synechocystis sp. PCC 6803. Biochim. Biophys. Acta Bioenerg. 1858, 510–518 (2017).

52. T. Malavath, I. Caspy, S. Y. Netzer-El, D. Klaiman, N. Nelson, Structure and function of wild-type and subunit-depleted photosystem I in *Synechocystis*. Biochim. Biophys. Acta Bioenerg. 1859, 645–654 (2018).

