## Supplemental Information for "Alka(e)nes contribute to membrane lipid homeostasis and resilience of photosynthesis to high light in cyanobacteria"

### Supplementary Information

Rui Miao[1,2]*, Bertrand Légeret[1], Stéphan Cuine[1], Adrien Burlacot[1,3], Peter Lindblad[2], Yonghua Li-Beisson[1], Fred Beisson[1], Gilles Peltier[1]*

[1]Institut de Biosciences et Biotechnologies Aix-Marseille, CEA CNRS Aix-Marseille University, CEA Cadarache, Saint-Paul-lez-Durance, F-13108 France
[2]Microbial chemistry, Department of Chemistry-Ångström Laboratory, Uppsala University, Box 523, SE-751 20 Uppsala, Sweden
[3]Carnegie Institution for Science, Department of Plant Biology, 260 Panama street, Stanford, CA 94305, USA

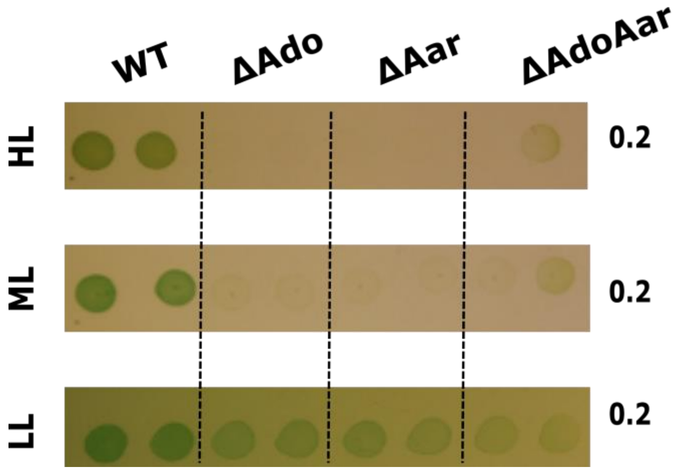


**Supplementary figure 1.** Growth test on BG-11 agar plate under different light intensities. Biological duplicates are shown for each strain. Initial chlorophyll concentration for each spot was 0.2 μg $ml^{-1}$. HL: 200 μmol photons $m^{-2}$ $s^{-1}$, ML: 50 μmol photons $m^{-2}$ $s^{-1}$, LL: 20 μmol photons $m^{-2}$ $s^{-1}$.

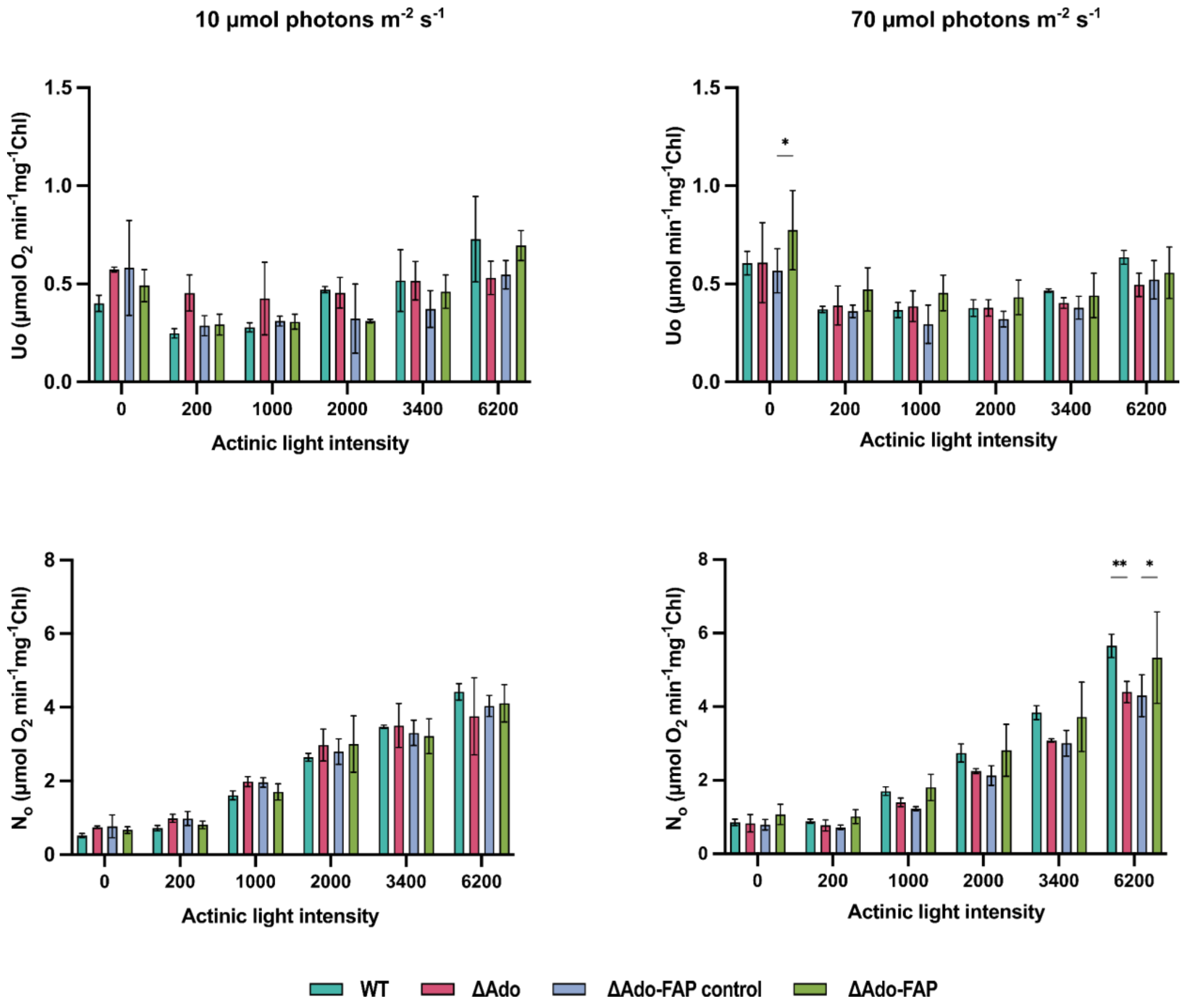


**Supplementary figure 2.** Membrane-inlet-mass-spectrometry (MIMS) measured $O_2$ uptake and net $O_2$ evolution rates from cells grown under low light and high light intensities. Measurements were done under increased blue actinic light. Results represent 3 biological replicates; error bars represent standard deviation, and significance was determined using two-way ANOVA.

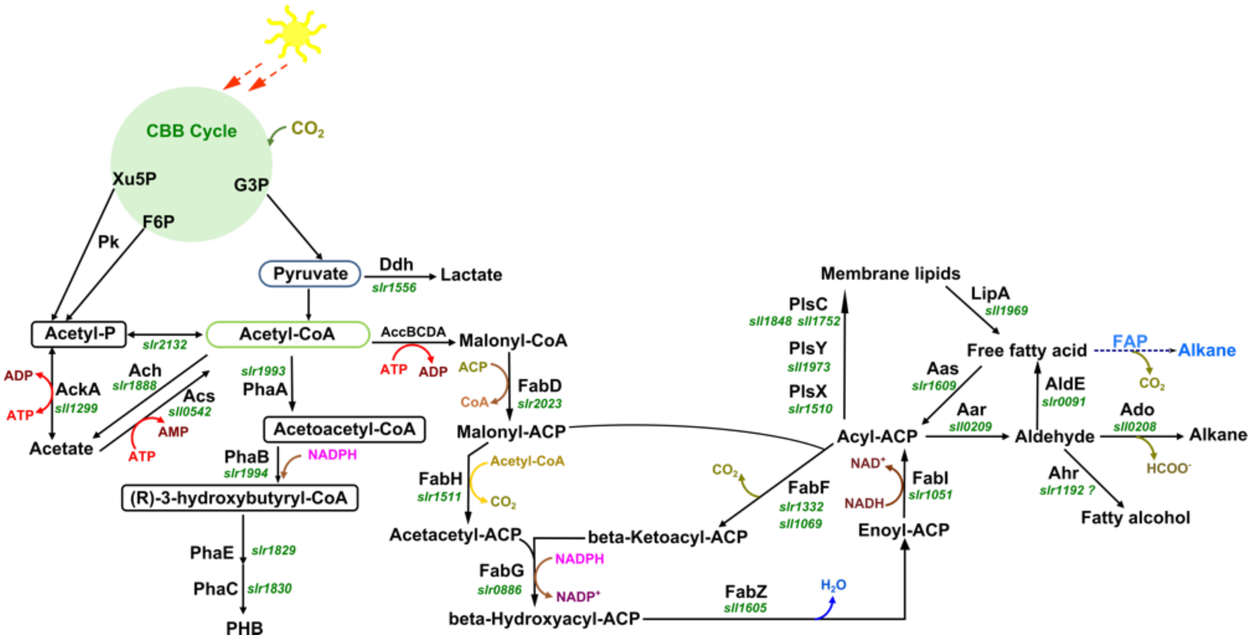


**Supplementary figure 3.** Scheme of central metabolic pathways relate to alka(e)nes synthesis in *Synechocystis* PCC6803

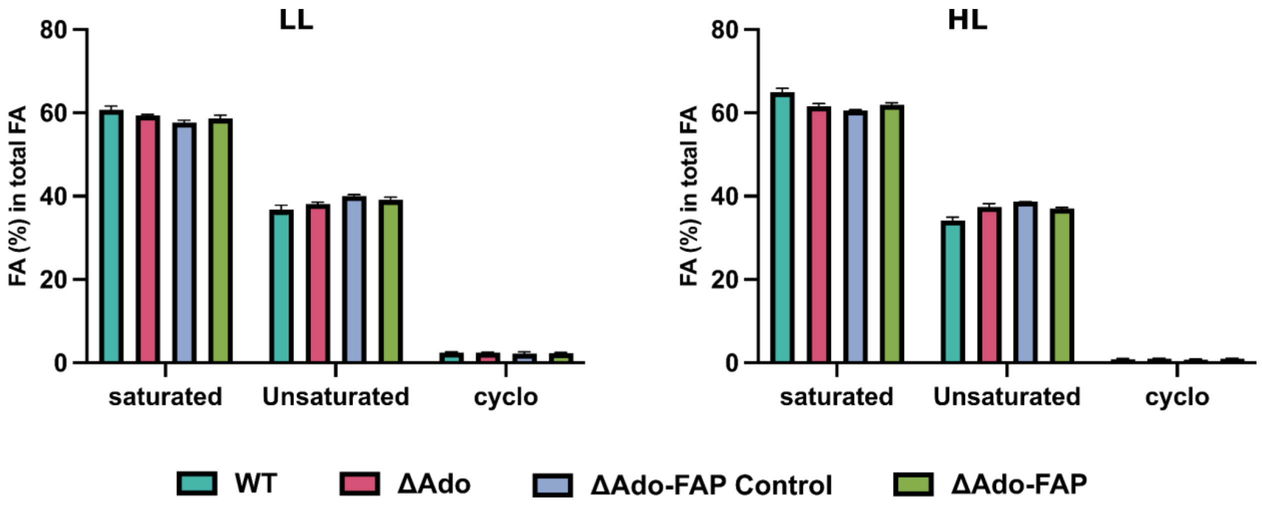


**Supplementary figure 4.** Proportion of saturated-, unsaturated-, and cyclo-fatty acid under low and high light. The content of each fatty acid species was normalized to the total fatty acid content of each sample, and the sum of saturated, unsaturated, and cyclo FA species contents was calculated. Results represent 3 biological replicates. Error bars indicate standard deviation, and significance was determined using a two-way ANOVA.

**A**

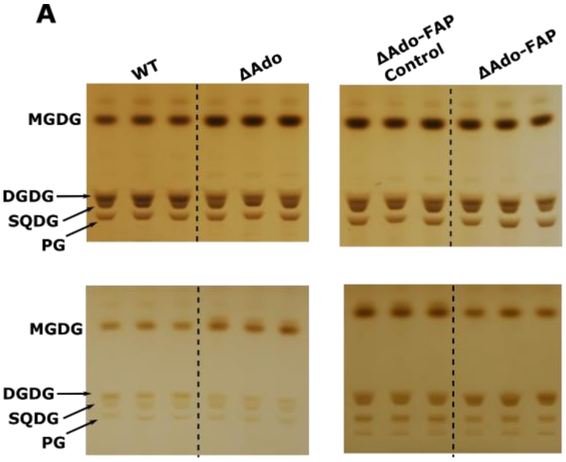


**B**

| | | SQDG% | PG% |
|---|---|---|---|
| Low Light | WT | 2.0±0.2 | 27.7±3.9 |
| | ΔAdo | 2.0±0.2 | 31.4±0.7 |
| | ΔAdo-FAP control | 3.4±0.6 | 36.9±4.9 |
| | ΔAdo-FAP | 2.6±0.9 | 33.4±3.1 |
| High Light | WT | 6.3±0.84 | 11.1±0.5 |
| | ΔAdo | 7.4±0.12 | 11.4±0.3 |
| | ΔAdo-FAP control | 6.9±0.86 | 11.6±0.13 |
| | ΔAdo-FAP | 6.7±0.19 | 14.9±2.77 |

**Supplementary figure 5. A.** Polar lipid quantification on High-Performance Thin Layer Chromatography (HPTLC). Upper panel: cells cultivated under high light. Lower panel: cells cultivated under low light. High-light and low-light cultivated samples were run as two separate sets of measurements. For each set, the same cell volume (representing biomass) was used for each sample. **B.** The proportion of SQDG and PG in total lipid content. Results represent the mean of 3 biological replicates; error bars represent standard deviation.

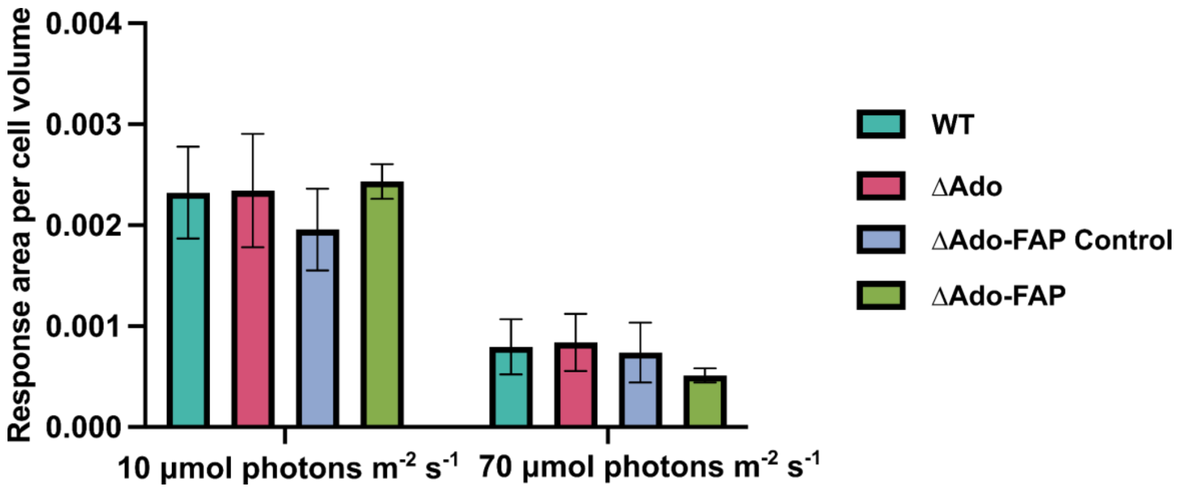


**Supplementary figure 6.** Chlorophyll content measured on Liquid Chromatography-Mass Spectrometry (LC-MS). Cells grown under low light and high light intensities were measured, and the response area on LC-MS was normalized by cell volume for each sample. Results represent 3 biological replicates; error bars represent standard deviation, and significance was determined using a two-way ANOVA.

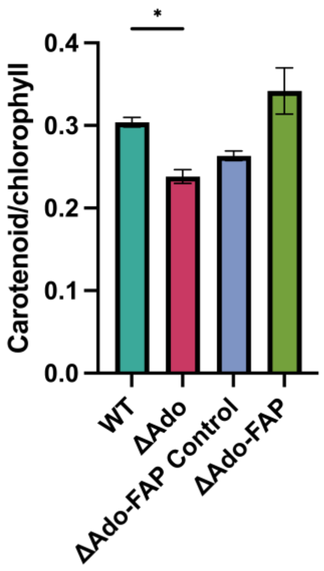


**Supplementary figure 7.** Spectrophotometer measured carotenoids to chlorophyll content ratio in cells cultivated under 70 μmol photons $m^{-2}$ $s^{-1}$ light with 1% $CO_2$. All results represent the mean of 3 biological replicates; error bars represent standard deviation, and significance was determined using one-way ANOVA.

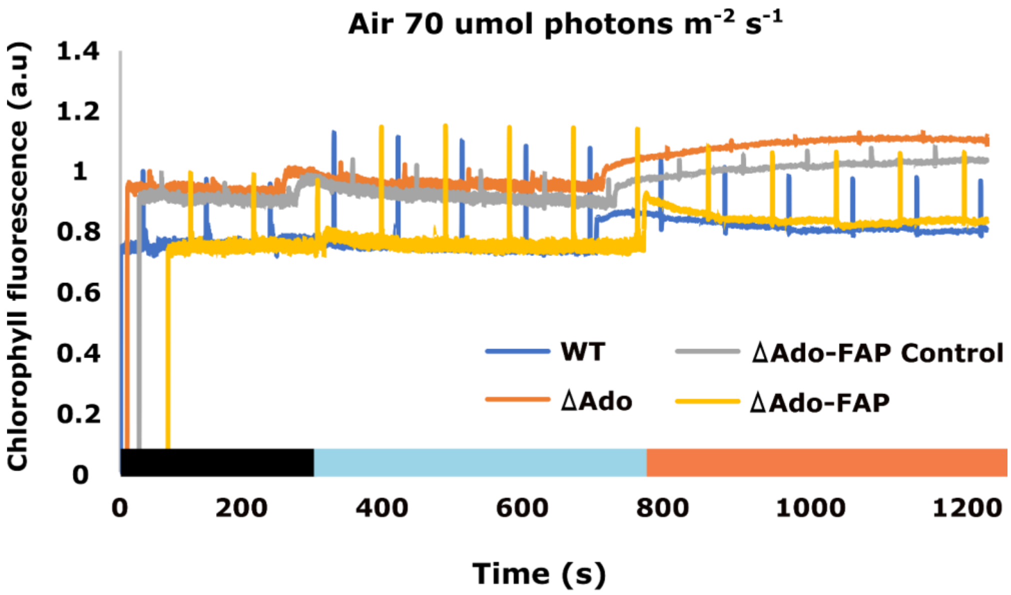


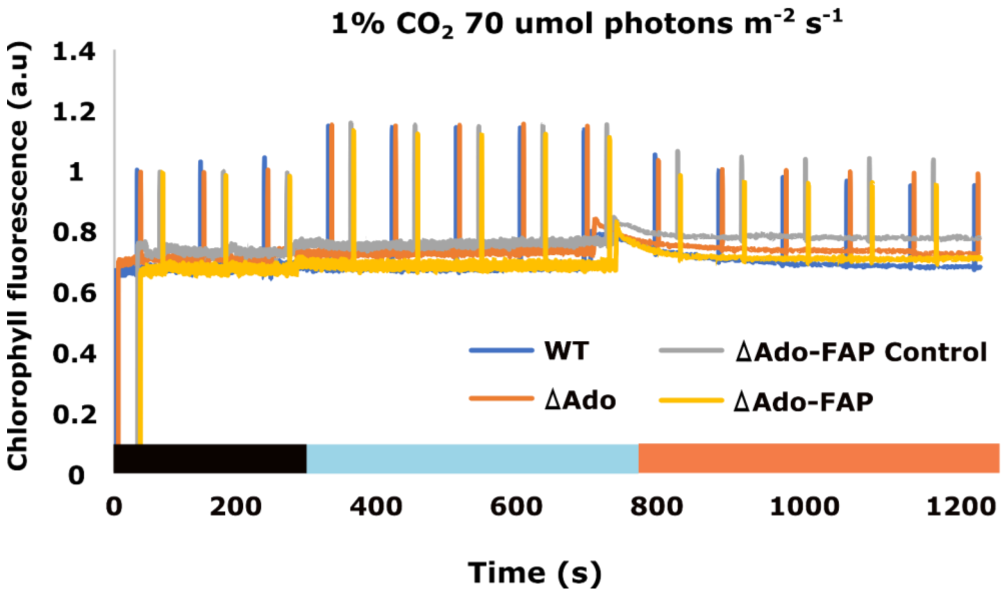


**Supplementary figure 8.** Chlorophyll fluorescence measured on cells grown in atmospheric level or 1% $CO_2$ conditions. Cells were cultivated under 70 µmol photons $m^{-2}$ $s^{-1}$, and chlorophyll fluorescence was measured in darkness, low blue actinic light (10 µmol photons $m^{-2}$ $s^{-1}$), and high orange actinic light (200 µmol photons $m^{-2}$ $s^{-1}$).

**Supplementary table 1.** List of engineered strains and genetic constructs

| Strain name | Genetic modification |
|---|---|
| ΔAdo | Δ*sll0208*::Cm$^{R}$ |
| ΔAar | Δ*sll0209*::Cm$^{R}$ |
| ΔAdoAar | Δ*sll0208-sll0209*::Cm$^{R}$ |
| ΔAdo-FAP Control | Δ*sll0208*::Cm$^{R}$ Δ*slr1993*::Sp$^{R}$ |
| ΔAar-FAP Control | Δ*sll0209*::Cm$^{R}$ Δ*slr1993*::Sp$^{R}$ |
| ΔAdoAar-FAP Control | Δ*sll0208-sll0209*::Cm$^{R}$ Δ*slr1993*::Sp$^{R}$ |
| ΔAdo-FAP | Δ*sll0208*::Cm$^{R}$ Δ*slr1993*::FAP-Sp$^{R}$ |
| ΔAar-FAP | Δ*sll0209*::Cm$^{R}$ Δ*slr1993*::FAP-Sp$^{R}$ |
| ΔAdoAar-FAP | Δ*sll0208-sll0209*::Cm$^{R}$ Δ*slr1993*::FAP-Sp$^{R}$ |
